# Whole-brain chemosensory responses of both *C. elegans* sexes

**DOI:** 10.1101/2025.05.15.654129

**Authors:** Maedeh Seyedolmohadesin, Xingyang Fu, Mahdi Torkashvand, Sina Rasouli, Samuel Lang, Liangzi Li, Cristine Kalinski, Steven J. Cook, Frank C. Schroeder, Eviatar Yemini, Vivek Venkatachalam

## Abstract

Sexually-dimorphic neural circuits play a critical role in shaping sex-specific animal behaviors. Maps of the structural dimorphisms in these circuits have been explored by analyzing “synaptic connectomes”, electron micrograph reconstructions of synaptic connectivity. Nevertheless, recent studies in the model organism *C. elegans* have shown little to no correlation between the synaptic connectome and dynamic neural activity. Therefore, the extent of sexual dimorphism in functional neural activity remains unknown. To determine the extent of functional sexual-dimorphisms in *C. elegans* we compared activity, neuron-by-neuron, across all neurons in the heads of both sexes. To sample a broad view of responses to different sensory modalities, we tested a diverse panel of ethologically-relevant olfactory, gustatory, and chemical stimuli, representing both attractive and aversive cues. We found that nearly every sensory neuron responded dimorphically to at least one cue and monomorphically to other cues, indicating that sexually-dimorphic circuits are pervasive and stimulus dependent. This dimorphic and monomorphic activity was present to a lesser extent in downstream interneurons and even less so in motoneurons, implicating sensory neurons as the primary source and location of sexually-dimorphic activity. Comparing the functional activity we measured to the published synaptic connectomes of both sexes revealed that sexual dimorphism in functional connectivity was distinct from and complementary to sexual dimorphism in synaptic connectivity. Our results provide a first-of-its-kind comparison of whole-brain dynamics between sexes at the level of single neurons, serving as an extensive resource for further investigations of functional sex differences.

## Introduction

During sexual maturation, animal nervous systems develop in a sexually differentiated manner to add critical reproductive behaviors. The resulting changes in neuronal function and connectivity lead to the generation of sexually-dimorphic behaviors, optimized for each sex’s reproductive role [1– 5]. Common examples of these include male courtship and female mate selection behaviors. Sexually-dimorphic behaviors are often activated by the presence of sensory cues. These sensory-driven dimorphic behaviors are especially exemplified by the pheromone responses of both invertebrates and vertebrates. Males of the model organism *Caenorhabditis elegans*, a millimeter-sized nematode, are attracted to pheromone released by the opposing sex [6]. Likewise, for fruit flies and mice, pheromones trigger male attraction to females and even aggression to other males [7–11]. More broadly, even sensory cues that trigger similar behaviors from both sexes can show functional sexual dimorphisms. For example, in *C. elegans* the opposing sexes use different neurons and circuits to respond to nociceptive stimuli [12–15]. Likewise, humans, non-human primates, and rodents show functional sexual dimorphisms in their nociceptive neurons [16]. Together these studies and others ([6, 8, 17–20]) demonstrate that sexually-dimorphic behaviors across species can be activated by dimorphic sensory perception.

Even when animals perceive their environment in a sexually monomorphic manner, their underlying nervous systems have been shown to employ sexually-dimorphic communication and processing. As a result, significant efforts have been dedicated to comparing neuronal morphology and connections between sexes across the nervous system. Connectomes for both sexes of the nematode *C. elegans* and the fruit fly *D. melanogaster* are now available and show a variety of sexual dimorphisms [21–27]. Comparative analysis in *C. elegans* has identified sex-specific neurons that are primarily responsible for behaviors unique to each sex, such as egg laying in hermaphrodites [21, 28] and copulation in males [29, 30]. Significant sexual dimorphism has also been observed between synaptic connections of neurons present in both sexes (sex-shared neurons), that contribute to dimorphic behaviors between the sexes [12, 23, 31]. Connectome studies in *D. melanogaster* have revealed fewer differences in the morphology and connections of neurons between the sexes despite their significant behavioral and transcriptomic differences. Specifically, in the fly optic lobe, only 0.2% of all cells are unmatched between males and females. Among descending and ascending neurons, 1-4% were found to be sex-specific, while 2-3% were sex-shared but with dimorphic morphology. Similar to *C. elegans*, these sexually dimorphic neurons are responsible for controlling sex-specific behaviors, such as egg-laying in females and courtship in males [24, 25]. In rodents, despite significant behavioral differences between the sexes, only a few anatomical dimorphisms have been observed [2, 18, 32].

Finally, even with identical sensory processing and synaptic connectivity, male and female nervous systems may process information in a dimorphic manner via (1) distinct intracellular computation, or (2) distinct intercellular nonsynaptic communication (i.e., volume transmission) via diffusive chemicals such as neuropeptides. These differences are evidenced by transcriptomic dimorphisms in molecules responsible for internal computation and external communication [33–40]. For example, in *C. elegans* 37 out of 154 neuropeptide genes are dimorphically expressed, and nearly all are upregulated in males during sexual maturation. Furthermore, conserved neuropeptides like oxytocin and vasopressin signal nosynaptically via volume transmission, acting sex-specifically in both mammals and nematodes [41–43]. This provides another clear pathway for substantial sexual dimorphism in neuronal computation.

Taken together, these advances demonstrate a clear need to characterize the complete extent to which a nervous system is functionally dimorphic and how closely this functional dimorphism aligns with structural, molecular and synaptic differences. The inability to evaluate functional dimorphisms represented a bottleneck for these comparisons. We recently developed the NeuroPAL method to simultaneously determine the neural activity and identity of every neuron in both sexes of *C. elegans* and thus evaluate functional activity, neuron-by-neuron, across the sexes [44, 45]. Here we leverage this advance to investigate neural activity and determine its relationship to synaptic connectivity in *C. elegans*, using NeuroPAL to measure functional dimorphisms and compare these with the connectomes of both sexes. [21–23].

In this study, we specifically asked: (1) How does the entire nervous system of each sex respond to a broad set of ethologically relevant sensory cues? (2) To what extent are individual neurons functionally dimorphic? and (3) How does the sexual dimorphism in functional connectivity correspond to dimorphism in synaptic connectivity? To address these questions, we developed a novel system to record neural activity across the nervous system (including both head and tail neurons) at single-cell resolution in both sexes of *C. elegans*, while presenting a diverse set of chemosensory cues to their nose. We characterized dimorphisms in individual neuron responses as well as in the functional connectivity between neuron pairs. Our findings revealed a high degree of dimorphism in sensory neurons and a low degree of dimorphisms in their functional connectivity (pairwise correlated activity), indicating that in *C. elegans* sexually dimorphic behaviors may be primarily mediated by dimorphism within sensory neurons. We additionally found that the observed dimorphisms in functional connectivity do not match dimorphisms in synaptic connectivity, suggesting that these dynamic and static circuit views represent complementary sex-specific communication pathways.

## Results

### Broad chemosensory stimulation activates the whole brain of both sexes

We investigated sexual differences in neural activity by measuring calcium activity, at single-neuron resolution, simultaneously for the head and tail of both sexes of *C. elegans* (Fig. 1). To broadly activate the head neurons we targeted diverse chemosensory circuits spanning a large variety of sensory, inter-, and motor neurons (Fig. 1C) [46]. We stimulated these overlapping circuits by presenting a diverse panel of ethologically relevant chemosensory stimuli to animal’s nose (Fig. 1D,E). For this stimulus panel, we selected 8 cues that sample a variety of modalities (Fig. 1C): (1) OP50 bacterial supernatant to represent food, (2) 1 µM ascr#3 to represent pheromones [47], (3) 800 mM sorbitol to represent osmotic stress [48], (4) 10 mM CuSO_4_ to represent toxic stress [49], (5) low (10^−6^ v/v) and (6) high (10^−2^ v/v) concentrations of isoamyl alcohol (IAA) a food-related olfactory cue [50, 51], as well as (7) low (100 mM) and (8) high (450 mM) NaCl to represent gustatory cues [52]. These stimuli evoked robust behavioral responses in both wild-type (N2) worms and the NeuroPAL;GCaMP6s (OH16230) worms which were used for imaging (Fig. 1C and S3). NeuroPAL strains employ an invariant multi-color, fluorescent-protein barcode to identify every neuron subtype in *C. elegans* of both sexes, across all developmental stages [44, 45]. NeuroPAL strains can be crossed to GFP/CFP/YFP reporters to identify gene expression, or neural-activity sensors such as GCaMP6s to measure neural activity [53]. The head of *C. elegans* are composed of 89 sex-shared neuron classes [21]. These highly-connected head neurons represent a primordial brain. Thus imaging their neural activity is called “whole-brain imaging” [54]. Presenting our stimulus panel to both hermaphrodites and males generated consistent whole-brain activity in 84 of 89 neuron classes, confirming broad activation of the head circuitry. Although head neurons exhibited broad activation in response to sensory cues, as seen in our prior work [44], tail neurons did not appear to be responsive to cues presented to the head. Therefore, we focused solely on analyzing the head neurons.

**Figure 1.**
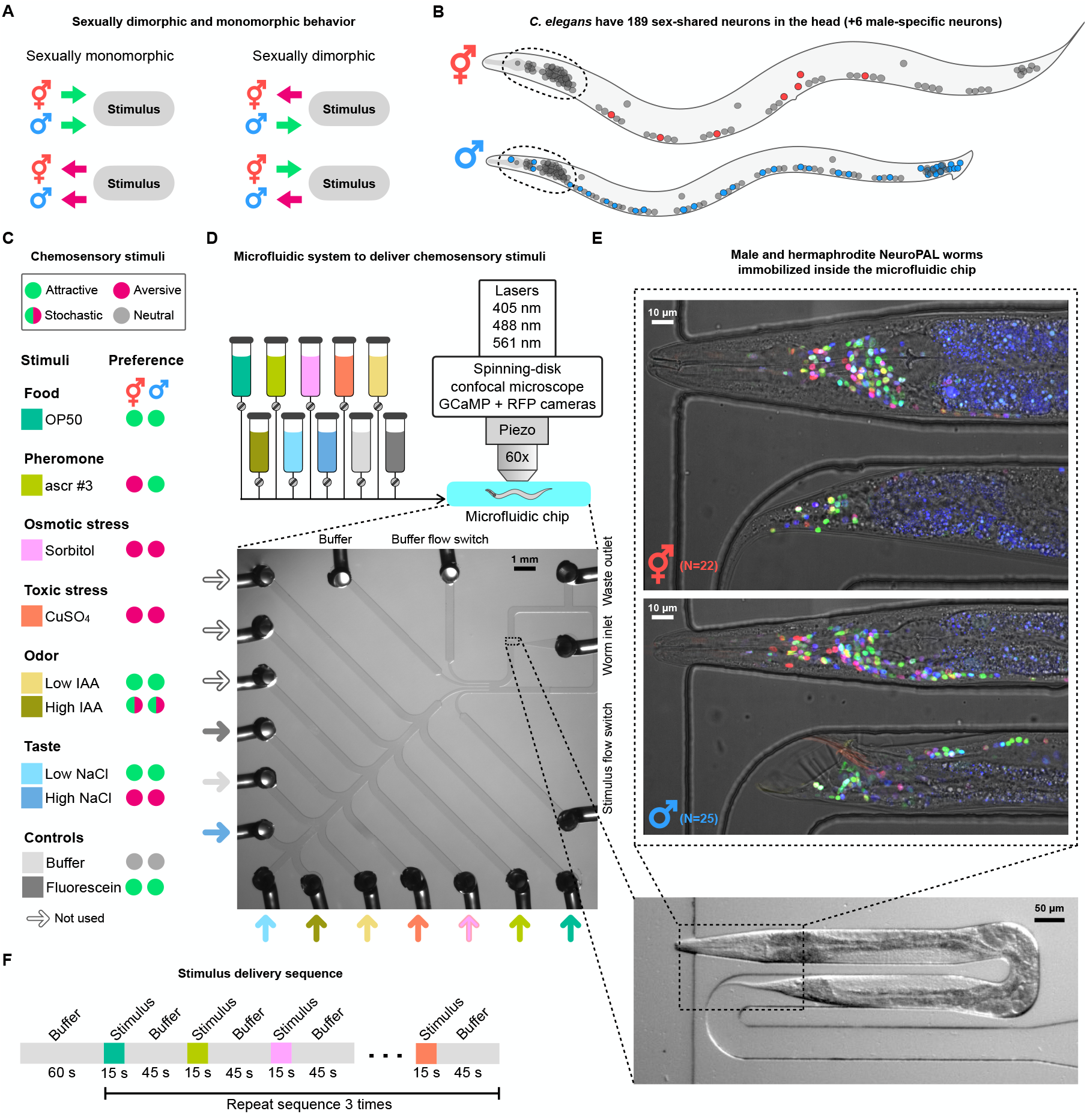
System for recording whole-brain activity of *C. elegans* in response to a large panel of stimuli in both sexes. **A**. Hermaphrodite and male *C. elegans* exhibit sexually monomorphic and dimorphic chemosensory preferences. **B**. Neurons in hermaphrodite (top) and male (bottom) *C. elegans*. Sex-specific neurons for hermaphrodites (red) and males (blue). The heads of both sexes contain 196 sex-shared neurons categorized into 89 classes. **C**. Eight stimuli that target multiple sensory modalities were used to activate behavioral circuits. Hermaphrodite and male preferences to these stimuli are shown in green (attractive), magenta (aversive), and gray (neutral). **D**. Diagram of the microfluidic imaging system, composed of a spinning-disk confocal microscope, cameras for fluorescent imaging, and a microfluidic device capable of serially delivering up to 13 stimuli to a single worm. **E**. NeuroPAL hermaphrodite (top) and male (bottom) immobilized inside the U-shaped trap of the microfluidic chip. **F**. Stimulus delivery sequence. Buffer was delivered for 60 seconds to acclimate animals to the 488 nm laser light. Stimuli were presented for 15 seconds, followed by 45 seconds of buffer. The order of stimulus presentation was randomly permuted for each animal, and repeated three times for that animal.

To sequentially deliver the stimuli listed above to each animal, we designed and fabricated a new microfluidic device that innovates on prior designs [44, 55, 56]. We designed two chips to accommodate the different lengths, widths, and depths of young-adult hermaphrodites and males (Fig. 1D,E). We used 13 stimulus channels to present aqueous chemicals to the nose of recorded animals [44, 56]. A U-shaped worm trap enforced lateral orientation [57] to capture both the head and tail of animals within the field of view of the microscope and cameras. This setup captured over 2/3 of the *C. elegans* nervous system, with a field of view (FOV) that encompassed approximately 230/302 neurons for hermaphrodites and 304/387 neurons for males; this FOV only excluded a subset of the ventral nerve cord and midbody neurons. We imaged the fluorescent proteins composing the NeuroPAL color barcode and the GCaMP6s neural-activity sensor using a spinning-disk confocal microscope (Fig. 1D), equipped with 3 laser lines (405, 488, and 561 nm) and filters for the NeuroPAL;GCaMP6s fluorescent proteins (mTagBFP2, CyOFP1, mNeptune2.5, TagRFP-T, and GCaMP6s). Neuronal calcium activity (GCaMP6s) was imaged at 2.67 volumes per second (25 frames per volume, 15 ms exposure per frame) using a 60x silicone objective with a vertical piezoelectric positioner. For each animal, we recorded neural activity for 31 minutes continuously (Fig. 1F). Because worms are sensitive to blue light [58], each animal was given 1 minute to acclimate to the 488 nm laser before presenting a series of chemical stimuli. After acclimation, each stimulus was delivered for 15 seconds, followed by 45 seconds of buffer as both a control and to permit neural activity to return to baseline. For each animal, we randomly permuted the stimulus order to avoid systematic sampling effects. We presented the selected random permutation of stimuli to that animal 3 times during the 31-minute recording.

To distinguish stimulus-driven neural-activity from artifacts and noise, we used two controls: (1) We used buffer as a neutral stimulus to control for responses elicited by non-chemical sources (e.g., mechanosensory or thermosensory stimuli caused by switching the flow of stimuli within the chip). (2) We used a low concentration of the fluorescent dye fluorescein (10 µM) to ensure the animal fit tightly within the microfluidic chip so that stimuli were only presented to its nose, not its body – whenever fluorescin was detected in the body chamber of the microfluidic chip, the experiment was ended and discarded.

### Most sensory neurons in the head exhibit sexually-dimorphic responses

We studied the activity of 196 sex-shared neurons and 6 male-specific neurons in the heads of 22 hermaphrodites and 25 males. An example recording from each sex is shown in Fig. 2A. We observed stimulus responses in sensory, inter, motor, and pharyngeal neurons at stimulus onset, offset, and sometimes both. Notably, many sensory neurons showed responses to nearly every stimulus, as shown in Fig. 2A, suggesting that stimulus encoding may require broad sensory activation. Of the 89 head neuron classes that we measured, 84 were clearly responsive to stimuli. Here, we define responsive as having at least one stimulus elicit a significantly different response as compared to the buffer-to-buffer control, at stimulus onset or offset, in at least one sex. Although a small set of neuron classes were classified as not clearly stimulus responsive, many of these still showed measurable calcium dynamics not synchronized with stimulus delivery. The male-specific head neuron classes (sensory neuron CEM and interneuron MCM) were predominantly unresponsive to stimuli and therefore were not investigated further (Fig. S3).

**Figure 2.**
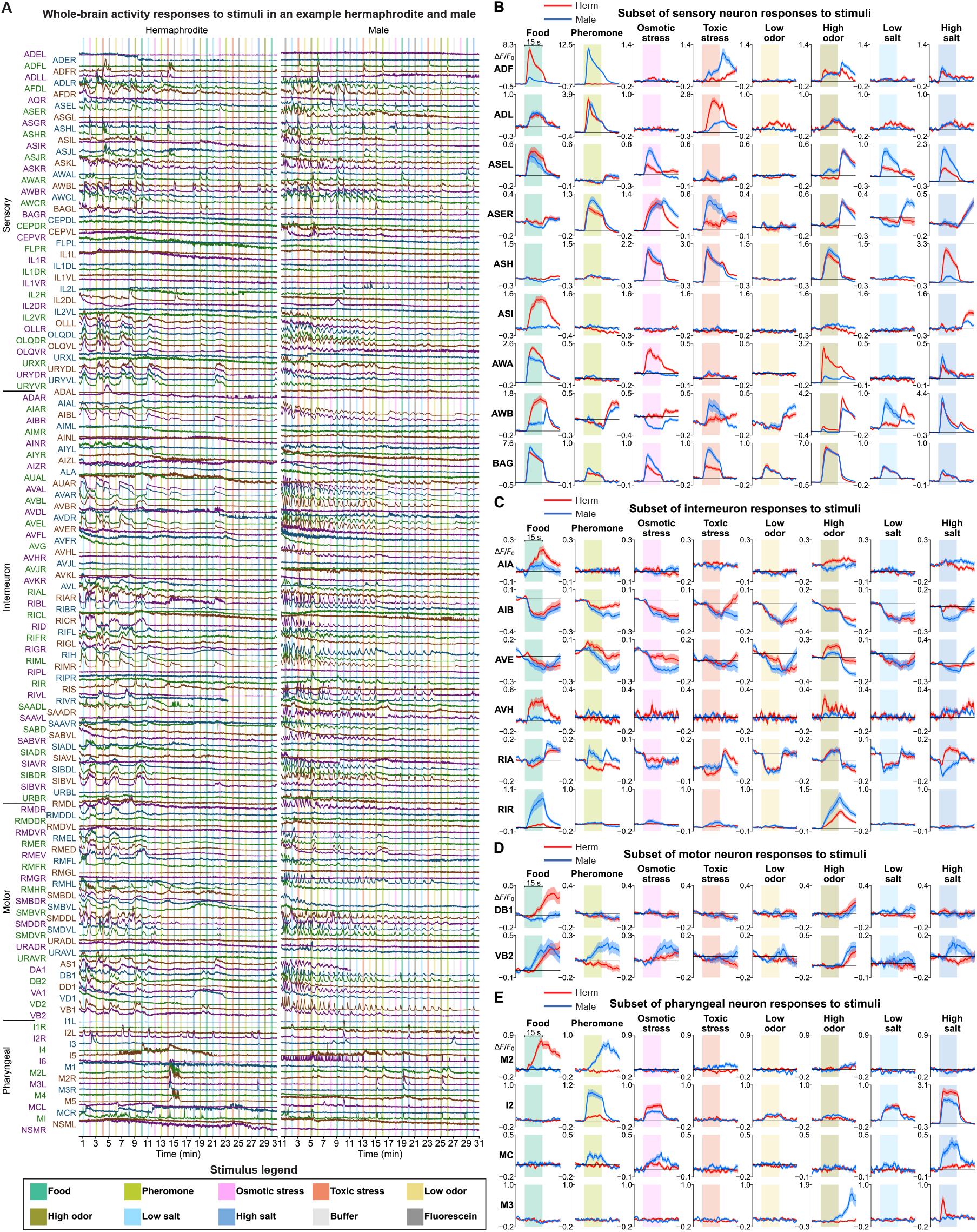
Whole-brain activity of both sexes in response to stimuli. **A**. An example of calcium activity in 196 sex-shared neurons from a hermaphrodite (left) and a male (right) in response to our panel of 8 stimuli and 2 controls. Each animal was continuously recorded for 31 minutes, during which the stimulus sequence was presented and repeated three times. To visualize the calcium traces we scaled them to have an equivalent range of fluorescence values (arbitrary units). Each stimulus was delivered for 15 seconds (colored vertical bars with color legend shown at the figure bottom). **B-E**. Mean hermaphrodite (red) and male (blue) stimulus responses for a subset of responsive neuron classes, exemplifying sexually dimorphic (e.g., BAG responding to high odor) and monomorphic neural activity (e.g., AWA responding to low salt). The mean activity (red and blue lines) with SEM (red and blue shaded boundaries) was calculated from 22 hermaphrodites and 25 males, combining neurons of equivalent left and right classes (e.g., combining nociceptive sensory neurons ASHL and ASHR), and computed using all 3 stimulus cycles. **B**. Sensory neurons. **C**. Interneurons. **D**. Motor neurons. **E**. Pharyngeal (enteric) neurons.

For both sexes, we calculated the mean response of each neuron class to each stimulus. *C. elegans* neurons are subdivided into classes that exhibit 2-fold symmetry (left / right), 4-fold symmetry (left / right and dorsal / ventral) or 6-fold symmetry (left / right, dorsal left / right, and ventral left / right). Neurons within the same class usually exhibit equivalent neural-activity traces. In this case, the convention is to combine their traces into a single class representation (e.g., combining activity from the left and right chemosensory AWA neurons, AWAL and AWAR, into a single AWA trace). The sole exceptions are the stereotypically asymmetric gustatory neurons ASEL and ASER [52] and the stochastically asymmetric olfactory neurons AWC-ON and AWC-OFF [59]. We did not observe large activity differences between AWC-ON and AWC-OFF, and occasionally had trouble distinguishing between them. For these reasons, we combined the mean activity of left/right neuron pairs into a single class representation across all three stimulation cycles, with the exception of ASEL/R which are represented independently (Fig. 2B-E).

We observed three characteristics in sexually-dimorphic stimulus responses: (1) dimorphism in the presence of a response at stimulus onset or offset, (2) dimorphism in the response amplitude, and (3) dimorphism in the response kinetics. These three qualitative differences in neural activity were generally well represented by a single statistic: the mean ∆*F/F*_0_ over a fixed stimulus-locked window. Despite the possibility that this statistic could fail to capture subtle dimorphisms which lead to similar means in neural activity, we did not find any such examples in our data.

For each stimulus tested, we observed at least one neuron class respond in a sexually dimorphic manner, and one class respond in a monomorphic manner. As an example, in response to the onset of osmotic stress, the gustatory neuron ASER and the nociceptive neuron ASH [60, 61] show monomorphic responses, whereas olfactory neurons AWA and AWB [62] show dimorphic responses (Fig. 2B and Fig. 3A). We further observed that many neuron classes can exhibit both sexually dimorphic and monomorphic responses depending on the stimulus. For example, the ASH neuron’s response to osmotic and toxic stress is monomorphic, but its response to high salt is dimorphic (Fig. 2B and Fig. 3A). Lastly, we found that single neuron classes can exhibit a monomorphic response to one concentration of a stimulus but a dimorphic response to another concentration. For example, the ASER neuron’s response to a low concentration of salt is dimorphic, but its response to a higher concentration of salt is monomorphic (Fig. 2B and Fig. 3A).

**Figure 3.**
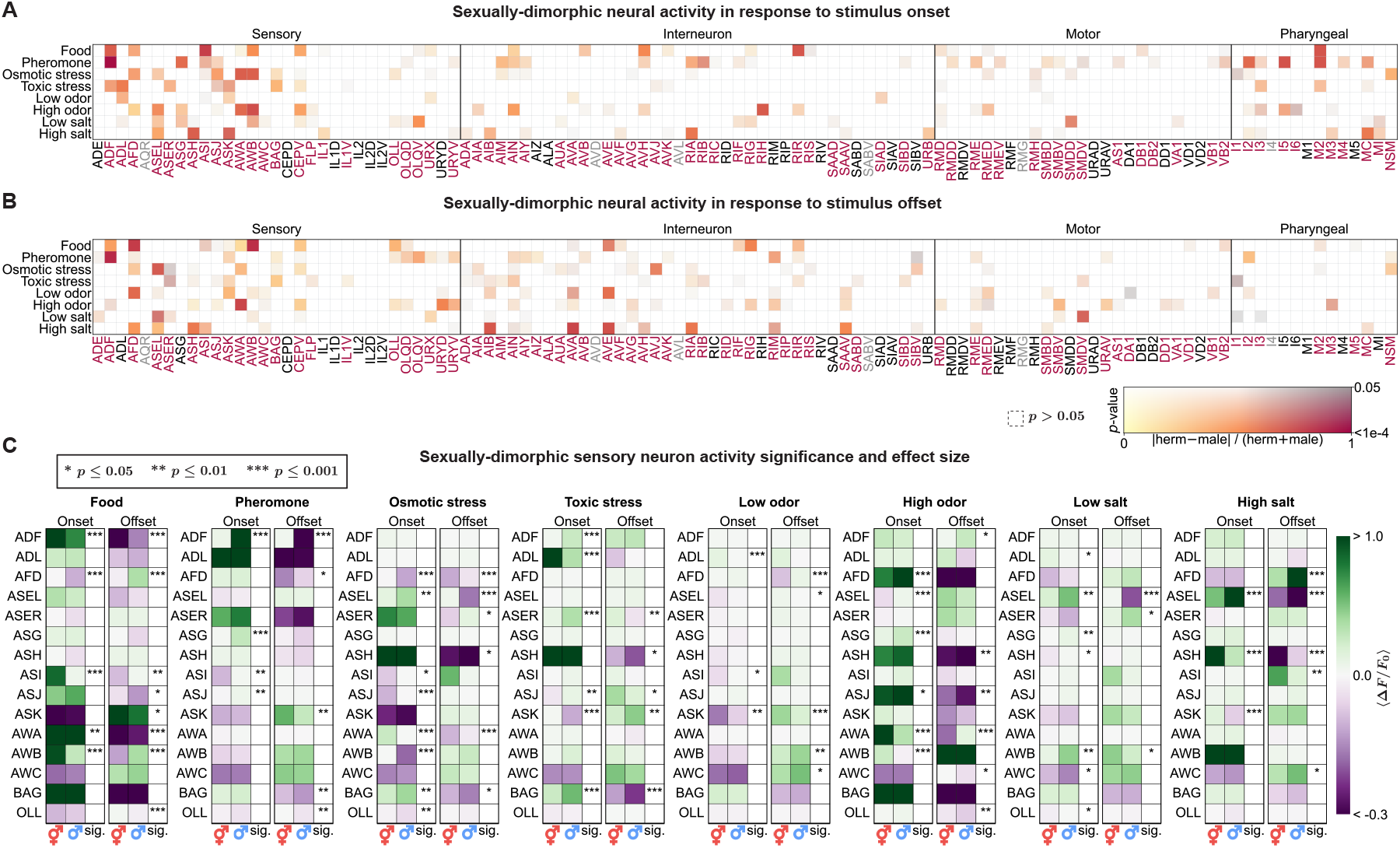
Sexual dimorphism in response to stimulus onset and offset. Quantification and significance of sexual dimorphism, per neuron subclass, for **A**. stimulus onset and **B**. offset. Stimulus onset and offset responses are computed as the mean calcium response during the 15 s period when the stimulus was presented, and during the 15 s period after the stimulus was stopped, respectively. Color indicates the relative sex differences in mean responses (red = large difference, yellow = no difference). Color brightness indicates the significance of the difference (bright colors = small more significant *p*-values, gray colors = large less significant *p*-values). Responses where *p* ≥ 0.05 are white, regardless of the relative difference (so that they can be ignored as insignificant). **C**. Comparison of mean responses (⟨∆*F/F*_0_ ⟩) for chemosensory neurons of both sexes for stimulus onset and offset. Stars indicate significance in male-hermaphrodite differences as determined by *p*-value (\**p* ≤ 0.05, \*\**p* ≤ 0.01, \*\*\**p* ≤ 0.001).

In addition to these dimorphisms, we noted several other surprising findings. Among the most remarkable of these were strong sexual dimorphisms in the pharyngeal neurons (Fig. 2B,E). The pharyngeal neurons represent a sub-network of 20 neurons that control feeding and that are connected to the main nervous system through a single gap junction between pharyngeal interneuron I1 and the head interneuron RIP. These neurons form a shallow network of inter- and motor-neurons with some sensory functions [63]. Therefore we were surprised to discover, for example, that pharyngeal interneuron I2 exhibited sexually dimorphic responses to pheromone, osmotic stress, and salt (Fig. 2E). Previous work had established that the I2 neurons regulate feeding in response to light [64]. Our discoveries suggest a far more substantial multi-sensory role for these enteric neurons that is further modulated by sex. In addition to this surprise, we found many examples of head neurons responding to their non-canonical stimuli. For example, as mentioned earlier, in addition to responding to odors, olfactory neurons AWA and AWB also responded to nociceptive stimuli including osmotic and toxic stress. Lastly, we observed that some neurons’ functions are switched between males and hermaphrodites. For example, ADF responds to food in hermaphrodites, but in males it responds to pheromones, matching expectations from [17]. More surprising was a similar dimorphic switch in the M2 pharyngeal neuron, also to food and pheromone (Fig. 2E). Taken together, these findings suggest that modifying sensory neuron functions may be a common and critical property in establishing sex-specific circuits.

We quantified the dimorphism of each neuron class in response to each stimulus for both stimulus onset and offset (Fig. 3A,B). To quantify the dimorphism of each neuron class, we first calculated the mean response over a 15-second period of stimulus delivery for stimulus onset and a 15-second period post-stimulus delivery for stimulus offset. We took the mean of these values across all individuals of each sex and across all three stimulation cycles. Sexual dimorphisms were then defined as the normalized difference between the mean hermaphrodite and mean male responses whenever this difference was statistically significant (*p*<0.05, independent sample t-test).

A significant fraction of sensory neurons responded dimorphically to at least one stimulus, and many responded dimorphically to multiple stimuli. Fig. 3C shows the mean calcium response (and its standard error) of each sex to each stimulus for both stimulus onset and offset. We observed that most neuron classes, across all neuronal layers, exhibited functional dimorphisms in their stimulus responses. Of the 89 neuron classes in the head, 84 were responsive (94%) and 80 classes (90%) were sexually dimorphic. Of the 24 sensory neuron classes, 23 classes (96%) were responsive and 22 classes (92%) were dimorphic. Of the 36 interneuron classes, 34 classes (94%) were responsive and 33 classes (92%) were dimorphic. Of the 15 motor neuron classes, 14 classes (93%) were responsive and 13 classes (87%) were dimorphic. Of the 14 pharyngeal neuron classes, 13 classes (93%) were responsive, and 12 classes (86%) were dimorphic (Fig. 5B).

All calcium activity data is available for both interactive exploration and downloading at https://chemosensory-data.worm.world.

### Functional correlations between sex-shared neurons are predominantly shared between sexes

The broad sexually-dimorphic responses that we observed may be due to either dimorphic responses to stimuli (e.g. dimorphic olfactory receptor expression), or dimorphic functional connectivity, in which signals propagate differently within the hermaphrodite and male nervous systems. To coarsely distinguish between these two possibilities we measured correlations between neurons and stimuli and between neuron pairs (functional connectivity) across the 31 minute recordings. To calculate the functional correlation between each neuron and each stimulus we encoded the stimuli as a binary-valued time series. To calculate the functional connectivity between different neuron subclasses representing all 110 L/R pairs and singleton neurons (Fig. 3), we calculated the mean Pearson correlation coefficient across all four possible combinations of neuron pairs (left-left, left-right, right-left, right-right), then computed the mean values across all individuals for each sex (Fig. 4A). This provided measures of stimulus correlation (880 neuron-stimulus pairs) and functional connectivity between each pair of neuron subclasses (5,995 pairs in each sex).

**Figure 4.**
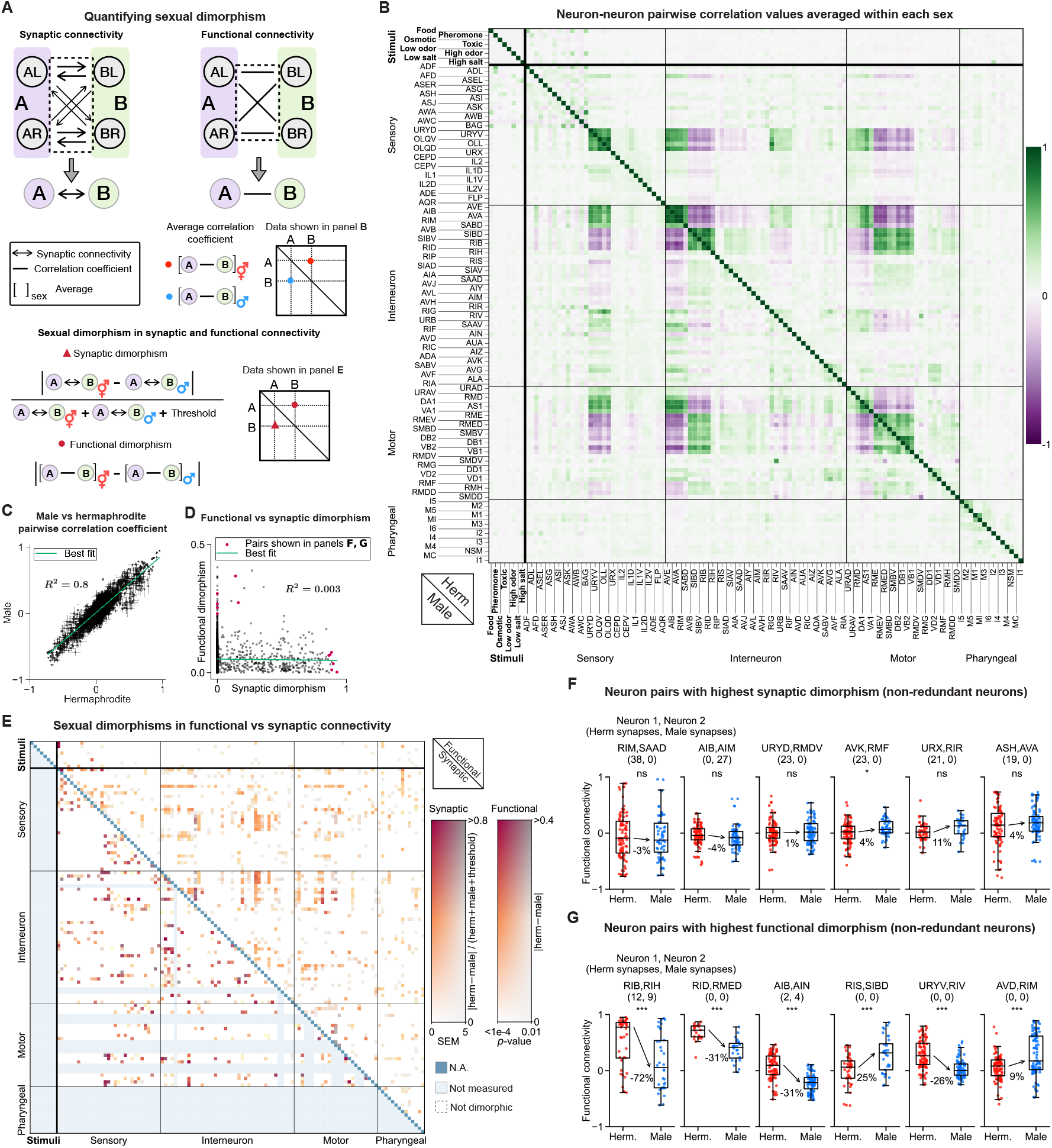
Comparing dimorphism in functional and synaptic connectivities. **A**. Illustration of methods to quantify sexual dimorphisms for functional and synaptic connectivity. The total number of synapses between neuron subclasses A and B is the sum of all 8 possible directional synapses. The correlation coefficient between neuron subclasses A and B is the computed as the mean Pearson correlation coefficient of traces from all 4 possible pairs. Synaptic dimorphism is calculated as the normalized difference in the number of synapses between sexes. Functional dimorphism is calculated as the absolute difference in the mean correlation coefficients between sexes. **B**. Neuron-neuron pairwise functional connectivity (magnitude of their correlation) shown as the mean across all hermaphrodites (upper diagonal) and males (lower diagonal), calculated as in panel A. **C**. Male and hermaphrodite functional connectivity is highly correlated (80%). **D**. Functional and synaptic dimorphisms are uncorrelated. Data at the extremes of functional and synaptic dimorphisms (red dots) is shown in panels F and G. **E**. Quantification of sexual dimorphism for functional (upper diagonal) and synaptic (lower diagonal) connectivity. Color indicates the numerical magnitude of functional and synaptic dimorphism (red = large difference, yellow = no difference). Color brightness indicates the statistical magnitude of the dimorphism. Functional dimorphisms are measured using *p*-values, and synaptic dimorphisms are measured using SEM due to low sampling: 3 hermaphrodite connectomes versus 1 male (bright colors = small more significant *p*-values or SEMs, gray colors = large less significant *p*-values or SEMs). Responses where *p* ≥ 0.01 or SEM > 5 are labeled white regardless of the dimorphism measured (so that they can be ignored as insignificant). We elaborate further on using SEM as a substitute for *p*-values in the main text. A version of this figure that includes the neuron names is available in Fig. S4. **F**. Neuron pairs with the highest synaptic dimorphisms (excluding redundancy in neural representation) show little to no dimorphism in functional connectivity. (\**p* ≤ 0.05, \*\**p* ≤ 0.01, \*\*\**p* ≤ 0.001) **G**. Neuron pairs with the highest functional dimorphism (excluding redundancy in neural representation), show little to no dimorphism in synaptic connectivity and are often not connected via synapses in either sex.

**Figure 5.**
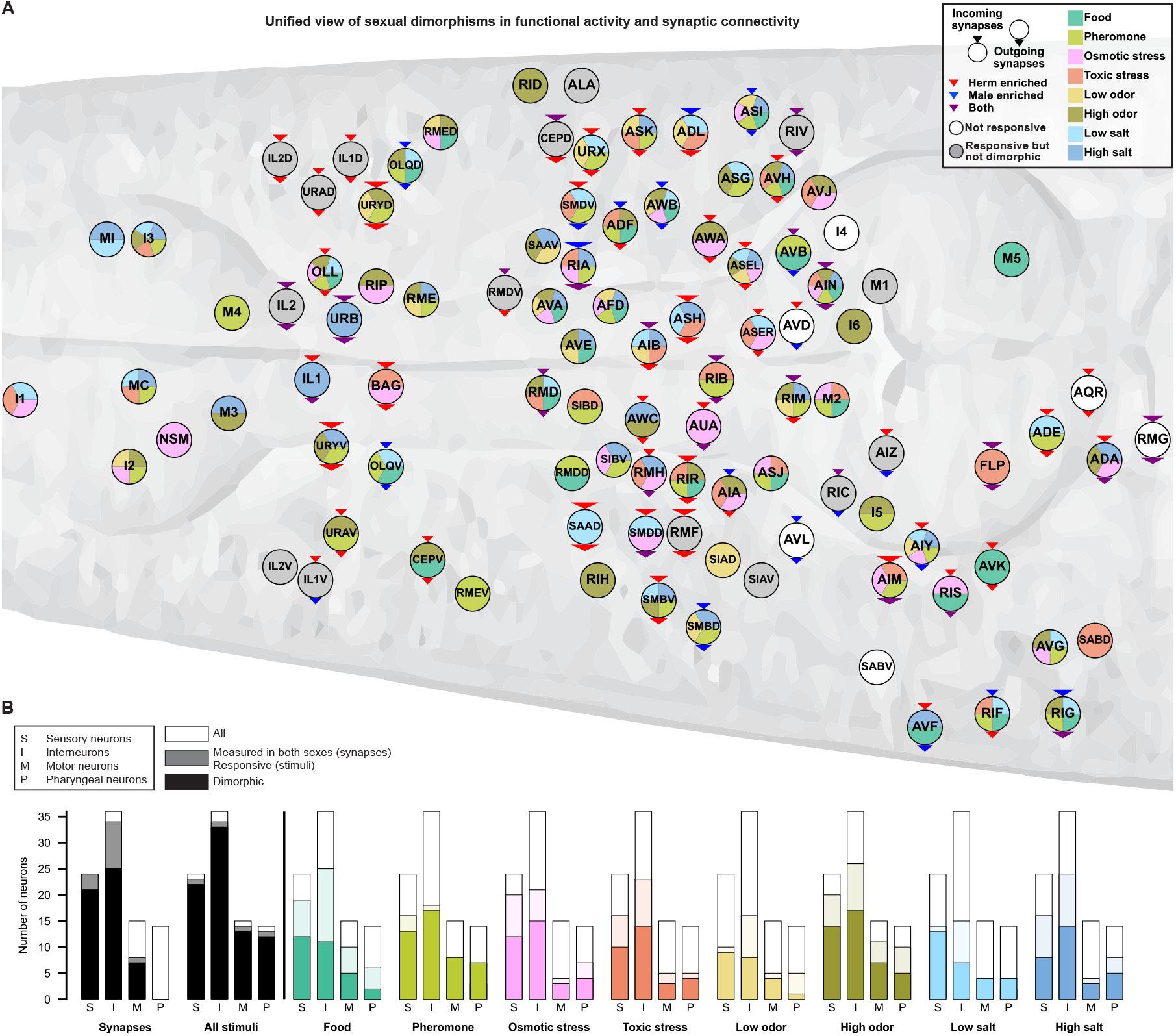
Sexual dimorphisms in functional activity and synaptic connectivity of each head neuron. **A**. Illustration of a *C. elegans* head and the neurons within, shown in their canonical positions. Each neuron is colored to show which stimuli, if any, evoked dimorphic responses. Dimorphic stimuli are shown as color-coded wedges. Neurons that were not responsive are shown in white. Neurons that were responsive but not dimorphic are shown in gray. Dimorphisms in incoming and outgoing synapses are shown as incoming and outgoing arrows whose size illustrates the magnitude of these dimorphisms. Red arrows signify that all dimorphic synapses are hermaphrodite-enriched. Blue arrows signify that all dimorphic synapses are male-enriched. Purple arrows signify that at least one dimorphic synapse is male-enriched and one is hermaphrodite-enriched. **B**. The number of dimorphic head neuron classes in each neuronal layer (sensory, inter, motor, and pharyngeal). In all plots, white indicates the total number of head neuron classes in each neuronal layer. In the first plot from the left, gray indicates the number of neuron classes where synapses were measured in both sexes, and black indicates the number of classes where either incoming or outgoing synapses were dimorphic. In the second plot from the left, gray indicates the number of neuron classes that were responsive to at least one stimulus, and black indicates the neuron classes that had a dimorphic response to at least one stimulus. In the remaining plots, light colors indicate the number of responsive neurons to that specific stimulus, and darker colors indicate the number of neurons that had a dimorphic response to that stimulus (same color legend as panel A). The classes used here are canonical *C. elegans* neuron classes (e.g., all six RMD neurons are collapsed into a single class). Fig. S6 provides a version with neuron subclasses to match Figs. 3 and 4.

The heatmap of the correlation coefficient values for both sexes is shown in Fig. 4B, with hermaphrodite values displayed above the diagonal and male values below the diagonal. We note that the heatmaps of both sexes are very similar. To quantify the similarity in functional connectivity between hermaphrodites and males, we plotted the mean pairwise correlation coefficient values for hermaphrodites versus males (Fig. 4C). This result shows that the functional connectivity between hermaphrodites and males is highly similar (80% or *R*^2^ = 0.8 for a linear fit). In both sexes, stimuli were more correlated (or anti-correlated) with sensory neurons than with interneurons or motor neurons. Within the nervous system, all pairs of neurons with strong correlation or anti-correlation show similar patterns in both sexes. Neurons within the pharynx show some correlations with each other but are largely uncorrelated with the rest of the brain, as expected for this synaptically isolated subnetwork [63].

### Functional and synaptic sexual dimorphisms are distinct

The whole-brain connectomes of both sexes [23] revealed dimorphisms in synaptic connectivity. However, we and others had previously shown that synaptic and functional connectivity are nearly entirely uncorrelated [44, 65–67]. Therefore, we investigated whether this lack of correlation between synaptic and functional connectivity extends to *sexual dimorphisms*. To measure synaptic dimorphisms between neuron classes, we summed all eight possible directed synaptic weights (Fig. 4A). We note that this metric does not discriminate between excitatory or inhibitory synapses, which are not differntiated in published connectomes. Two additional hermaphrodite connectomes were recently published [68]. Therefore, we compared the three available hermaphrodite connectomes to the single one available for males. We calculated dimorphisms in synaptic strength as the normalized difference between hermaphrodite and male synaptic strengths for each neuron pair (Fig. 4A). This resulted in a value between 0 and 1, with 1 indicating the highest dimorphism and 0 indicating no dimorphism. To estimate variability in synaptic strength for a given pair of neurons, we used the standard error of the mean (SEM) across the three hermaphrodite connectomes. Only one male connectome exists, so we were unable to do the same for males. This SEM was used as a reference to determine whether synaptic dimorphisms were significant or not.

Dimorphism in functional connectivity was defined as the absolute difference between the male and hermaphrodite mean correlation coefficients for each neuron pair. This resulted in a value between 0 and 2, with 2 indicating the highest dimorphism and 0 indicating no dimorphism. Despite this range, we observed that almost all neuronal pairs had a functional dimorphism of less than 0.4. These small differences in functional dimorphism highlight the overall similarity in functional connectivity between the sexes. To evaluate the significance of these functional dimorphisms, we performed an independent samples t-test between hermaphrodites and males for each pair of neurons, then used the resulting *p*-values as a measure of significance.

Fig. 4E shows the heatmap of dimorphism in functional connectivity versus that of synaptic connectivity. The two heatmaps display strikingly different patterns of dimorphism. We found no clear relationship between synaptic and functional dimorphism (Fig. 4D). To investigate this relationship in greater detail, we examined neuron pairs with the highest synaptic dimorphisms (Fig. S5A). These pairs were functionally uncorrelated in both sexes Fig. 4D. To visualize these differences, we show a subset of neurons with the highest synaptic dimorphisms in Fig. 4F. We further examined neuron pairs with the highest functional dimorphisms (Fig. S5B). We found that these pairs were mostly not synaptically connected. When they were connected, the connectivity was essentially not dimorphic. To visualize these differences, we show a subset of neurons with the highest functional dimorphisms in Fig. 4G.

## Discussion

Here we present, a first-of-its-kind resource that compares whole-brain sexual dimorphisms at a neuron-by-neuron level across the heads of opposing sexes. This resource includes both spontaneous neural activity as well as evoked responses to a broad panel of chemical cues spanning different sensory circuits and behavioral valences (Fig. 5). Our broad survey of sensory modalities and valences reveals that nearly every sensory neuron responds in a sexually dimorphic manner to at least one stimulus while responding monomorphically to at least one other stimulus. This suggests that a large fraction of dimorphic behavior in the worm may be explained primarily by cue-specific remodeling of the sensory layer that then feeds into mostly sex-shared circuitry downstream. This is consistent with previous reports of sexually-dimorphic sensory neuron classes in *C. elegans* [6, 12–14, 17, 22, 34, 47, 69–71], and prior work that suggests the inter- and motor-neurons are not dimorphic [72].

We also made the surprising discovery that the enteric nervous system of the worm responds to food, salt, and (most strikingly) pheromone in a sexually dimorphic manner (Fig. 2E). This enteric system is an isolated network of 20 pharyngeal neurons connected to the primary nervous system via a single gap junction (between neuron classes I1 and RIP) [21]. These enteric neurons have been primarily characterized as polymodal combinations of sensory, inter-, and motor neuron function [63]. While it is possible that these pharyngeal responses are downstream of this single gap junction bottleneck, we believe the richness of responses suggests that the pharynx is sensing chemicals directly in a dimorphic matter (Fig. 3A). The unexpected sexual dimorphism we find in this foregut suggests that sex may guide dimorphic responses related to feeding and metabolism via as yet uncharacterized pathways.

Functional correlations between neurons relates to their communication [73, 74]. Prior work evaluated synaptic dimorphisms using the connectomes of a single hermaphrodite and male [23]. We have updated this comparison by leveraging two more recently published connectomes for the hermaphrodite nerve ring, resulting in a comparison of three hermaphrodites and one male [68]. Published comparisons between functional activity and synaptic connectivity [44, 65–67, 75] have identified little to no simple correlation between the two, suggesting the relationship between calcium activity and synaptic structure is complex. Interestingly, we found that this extends to the relationship between sexual dimorphisms in functional activity and synaptic connectivity. Neurons with strongly dimorphic functional activity typically show no dimorphisms in synaptic connectivity between sexes (i.e., identical synaptic counts across both sexes), and vice versa (Fig. 4). Our findings indicate that functional activity and synaptic connectivity represent complementary views of how sexual identity is represented in *C. elegans*. Relating these views, which capture features of their behavioral circuits at different temporal and spatial scales, remains an outstanding challenge.

Though we focused on the brain of *C. elegans*, our observations may capture broad principles of how animals add sex-specific behavioral circuits during maturation. Notably, previous *C. elegans* and invertebrate studies of sexual dimorphisms have illustrated general mechanisms for sex-specific circuit assembly [23, 76–80] The overall and cue-specific functional sexual dimorphisms, along with our updated measurements of sexually dimorphic synaptic connections, across the brain of *C. elegans* are summarized in Fig. 5. This summary and our large accompanying resource illustrate basic fundamental mechanisms of how multimodal cues are dynamically encoded and processed in the brains of opposing sexes.

## Materials and Methods

### Worm maintenance

Worms were maintained at 20°C on Nematode Growth Medium (NGM) agar plates seeded with *E. coli* OP50, following published conventions [81]. Males were generated using standard heat-shock protocols, then backcrossed to hermaphrodites of the same strain to maintain a constant stock of isogenic hermaphrodites and males [82]. For all experiments, we used males that were passaged for at least four generations after the initial heat-shocking protocol.

### Worm Strains

All experiments were conducted with OH16230 hermaphrodites and males (otIs670[NeuroPAL]; otIs672[panneuronal::GCaMP6s]) [44].

### Behavioral assays

For stimulus drop tests, we used standard dry drop tests with CTX buffer [83, 84].

### Microfluidic chip design

The microfluidic chip was designed using elements of the olfactory chips described in [55, 56]. The design includes 2 switch channels, 1 buffer channel, and 1 main stimulus channel that splits into 13 different channels. At any given time, buffer, stimulus, and either of the 2 switches are active. These switches control whether buffer or stimulus flows in front of the worm. Switching from one stimulus to another occurs in the background during buffer delivery. The switch timing was set to ensure the new stimulus runs in the background for about 40 seconds before being delivered to the worm, allowing sufficient time for the removal of residues from the previous stimulus. The dimensions of the worm trap differed between males and hermaphrodites to account for morphological differences. To ensure that the rest of the chip remained identical between these two versions, we separated the chip into 2 layers: a worm trap layer and a stimulus/buffer flow-channel layer. The height of the flow channels is 70 *µ*m. The hermaphrodite trap has a width of 55 *µ*m and a depth of 28 *µ*m. The male trap has a width of 41 *µ*m and a depth of 25 *µ*m.

### Microfluidic chip fabrication

We designed the chips using AutoCAD software and sent the designs to CAD/Art Services, Inc. (now Artnet Pro, Inc.) for photomask printing. Standard photolithography was performed in a clean room to fabricate silicon wafer molds using the negative photoresist SU-8 2025. Polydimethylsiloxane (PDMS) was then poured over the mold and cured on a 90°C hotplate for 30 minutes to solidify. Individual chips were cut with a scalpel, and inlet and outlet holes were punched with a 1 mm biopsy punch. The chips were then bonded to No. 1.5 coverslips using a handheld plasma treater.

### Reagents and Stimulus Preparation

We used standard CTX buffer (5 mM KH_2_PO_4_/K_2_HPO_4_ at pH 6, 1 mM CaCl_2_, 1 mM MgSO_4_, and 50 mM NaCl) [85] as the solvent or diluent for all the stimuli except for CuSO_4_. For the food stimulus, we grew OP50 in LB overnight and resuspended it in CTX buffer one day before the experiment. 10 *µ*M stock solutions of ascr#3 in diH_2_O were stored in a -20°C freezer and were thawed and diluted in CTX to the final concentration of 1 *µ*M on the day of the experiment. Pure isoamyl alcohol was diluted in CTX on the day of the experiment. CuSO_4_ did not dissolve in CTX buffer, so we dissolved it in diH_2_O containing 50 mM NaCl. To minimize contamination, each stimulus had its own glass syringe and tubing for use in the microfluidic system.

### Preparation of worms for imaging

To ensure that mating did not occur prior to imaging, L4 stage hermaphrodites and males were separated the evening before the experiment and transferred to separate NGM plates that were seeded with OP50 bacteria for feeding. Worms were partially immobilized and loaded into the microfluidic device using 1 mM levamisole disolved in CTX buffer; with this concentration of levamisole, small movements were still visible.

### Multi-channel imaging and neuron identification

At the start of each experiment multi-channel imaging was performed to capture fluorophores expressed in NeuroPAL transgenic animals. The fluorophores imaged were mTagBFP2 (blue landmarks), CyOFP (green landmarks), mNeptune2.5 (red landmarks), TagRFP-T (a panneuronal marker), and GCaMP6s (a panneuronal calcium sensor). mTagBFP2 was imaged using a 405 nm laser and a Chroma ET450/50m emission filter. CyOFP was imaged using a 488 nm laser and a Chroma ET600/50m emission filter. mNeptune2.5 was imaged using a 561 nm laser and a Chroma ET700/75m emission filter. TagRFP-T was imaged using a 561 nm laser and a Chroma ET600/50m emission filter. GCaMP6s was imaged using a 488 nm laser line and a Chroma ET525/50m emission filter. Neuron identification was performed manually post-hoc after data collection as previously described [44, 45].

### Calcium imaging

Whole-brain activity was measured using the calcium sensor GCaMP6s by continuously recording for 31 minutes at cellular resolution using a custom spinning-disk confocal system similar to [86] with custom software. A detailed description is available at https://github.com/venkatachalamlab/lambda. Briefly, imaging was done with a piezo-driven Olympus 60x silicone immersion objective and Andor Zyla 4.2p sCMOS cameras attached to an Andor Dragonfly 200 spinning disc confocal with the laser lines, dichroics, and emission filters listed above. We recorded activity at 2.67 volumes per second. Volumes consisted of 25 frames, each exposed for 15 ms. Volume dimensions were 205 µm x 205 µm x 30 µm (x, y, z).

### Stimulus delivery

The start of the microfluidic experiment was synchronized with calcium imaging to ensure the exact same timepoints of stimulus onset and offset for each animal. To avoid bias from stimulus order, for each animal a randomized permutation of 10 stimuli was delivered, with the sequence repeated three times. Buffer was delivered for 60 seconds at the start of the experiment. Thereafter, each stimulus was presented for 15 seconds, with a 45-second buffer period between stimuli. We used a buffer-to-buffer stimulus control, delivered through a stimulus channel, to measure potential artifacts from channel switching (e.g., mechanosensory feedback or thermosensory differentials). We used this control to ensure neural activity responses were due to the stimuli presented and not artifactual. In each stimulus cycle, we used a fluorescein stimulus flow to ensure the microfluidic device was functioning and that the worm’s nose formed a sufficiently tight seal to prevent stimuli leaking into the body chamber (i.e., to ensure only the worm’s nose sensed our stimuli). If fluorescein leaked into the body chamber, we discarded the experiment. Air bubbles can occasionally flow into the chip from the microfluidic tubing and block or alter the flow direction, resulting in unreliable stimulus or buffer delivery. Therefore, we used fluorescein to ensure that stimulus delivery was not being impacted by potential air bubbles.

### Neuron tracking

Animals were immobilized using a combination of a nearly exact fit within the microfluidic chamber and anesthetic (1 mM levamisole). This allowed the animals to move slightly and contract their pharynx. We used ZephIR, a multi-object tracking algorithm described in [87], to track neuron centers at all time points. For each neuron, we calculated the calcium activity at each time point by taking the average intensity of the 15 brightest pixels within a rectangular volume of size 2 µm × 2 µm × 3 µm (x, y, z), centered in the middle of the neuron as determined by the tracking algorithm. Sample traces are shown in Fig. 2 A.

### Calculating stimulus onset and offset responses

To calculate a neuron’s response to stimulus onset and offset, we calculated ∆*F/F*_0_, where *F*_0_ is the average intensity over a 10 frame window (3.75 seconds) before stimulus onset/offset. To determine the average neuron response for each sex, across all trials and individuals, we excluded responses that were identified as substantial outliers using the statistical method described below. For each response, we first calculated the mean calcium activity (⟨∆*F/F*_0_ ⟩ _*t*_) during three time intervals, each with a duration of 15 seconds: 1. Before stimulus onset, 2. During stimulus delivery, 3. After stimulus offset. For each of these intervals, we pooled the data from all three stimulus cycles, across all individuals within each sex, and by combining neurons within each class. We then calculated the following: *IQR* = *Q*3− *Q*1, where *Q*1 is the 25th percentile and *Q*3 is the 75th percentile of the pooled data. We defined the lower threshold as: *Q*1 −5 · *IQR* and the upper threshold as: *Q*3 + 5 · *IQR* Data points below the lower threshold or above the upper threshold were identified as substantial outliers and subsequently excluded from further analysis. These outliers typically arose from errors tracking neurons between frames.

### Identifying responsive neuron classes

We identified neuron classes as responsive in at least one sex. A neuron class was marked responsive if its response to at least one stimulus was significantly different than the control response (buffer-to-buffer switch) at stimulus onset or offset. Significance was tested with the null hypothesis that the mean calcium response (⟨ ∆*F/F*_0 *t*_ *⟩*) to the stimulus was the same as the response to the control. For a given neuron class, we averaged across all animals of a given sex, across 3 stimulus cycles, and across left/right pairs (if applicable). Then we calculated the difference between the mean calcium responses to the stimulus and control. Neuron classes were deemed responsive if the *p*-value was ≤ 0.05, in at least one sex, in response to at least one stimulus, at either stimulus onset or offset.

### Identifying sexually dimorphic responses

We used a similar approach to identify sexually dimorphic responses. Significance was tested with the null hypothesis that the mean calcium response (⟨∆*F/F*_0_ ⟩ _*t*_) to the stimulus was the same in hermaphrodites and males. For every odor, neuron class, and stimulus timing (onset or offset) we calculated the difference between the mean calcium responses in hermaphrodites and males. Neuron classes were deemed sexually dimorphic if the *p*-value was ≤ 0.05 in response to at least one stimulus, at either stimulus onset or offset.

### Calculating dimorphism in functional connectivity

All neural calcium traces decay due to bleaching, which can lead to artifactual positive correlation coefficients between neuron pairs. To correct calcium traces for this bleaching, we band-pass filtered traces. For the band-pass filter, we subtracted the average signal in the past 240 frames (90.0s) from the average in the past 10 frames (3.75s) to restrict attention to the 11 mHz to 267 mHz range. After this filtering, we calculated the Pearson correlation coefficient for each pair of neurons across the entire recording. Each pairwise correlation was then averaged across all individuals of a given sex. We then quantified sexual dimorphism in functional connectivity by calculating the absolute difference between averaged hermaphrodite and male correlations. We identified significant dimorphisms using an independent t-test with *α* = 0.05.

### Normalization of connectome data

In this study, we used connectome data from four animals: 3 hermaphrodites and 1 male [21–23, 68]. These datasets were collected and analyzed by different groups at different times. To normalize the hermaphrodite data to accommodate these discrepancies, we performed the following computations. For each directed synapse between two neurons, we calculated the standard error of the mean (SEM) across the 3 hermaphrodite datasets. We then calculated the average and standard deviation of the SEM values across all synapses. This average SEM was 1.14 with a standard deviation of 2.07. To reduce the average SEM, we normalized each dataset by its 95% quantile value. This normalization resulted in synaptic weights that were mostly less than one. Therefore, we scaled all values by a fixed factor, specifically the 95% quantile of one of the hermaphrodite datasets (from [21]) which acted as the template for the remaining datasets. This method resulted in approximately a 50% reduction in the SEM values. Specifically, the average SEM decreased to 0.62 (down from the initial 1.14), and the standard deviation of the SEM decreased to 0.87 (down from the original value of 2.07). For our analyses, we represented the hermaphrodite synaptic weights as the average of their 3 normalized values along with the SEM.

### Calculating dimorphism in synaptic connectivity

Functional connectivity correlations are not directional. Therefore, to compare functional with synaptic connectivity we calculated the undirected synaptic strength between pairs of neurons. To do so, we summed all 8 possible directed synaptic weights (2 directions × 4 L/R combinations) for each sex. For hermaphrodites, we used the average synaptic weights across all 3 hermaphrodite datasets. As a measure of uncertainty in the combined synaptic strength, we applied error propagation rules to the SEM values of the individual synaptic weights, yielding the combined SEM for the sum:

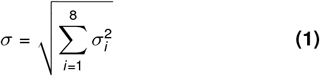

where *σ*_*i*_ represents the SEM of each individual synapse, and *σ* is the combined SEM.

To quantify dimorphism in synaptic connectivity, we calculated the following metric for each pair of neurons:

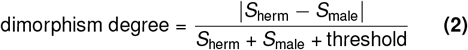

where *S* denotes the combined synaptic weight. The threshold value was included for regularization to ensure that synapses with very small synaptic weights do not exhibit disproportionately large degrees of dimorphism. This threshold value was set to 3. In other words, the difference between hermaphrodite and male synaptic weights must be at least 3 to qualify as a substantial dimorphism.

### Selecting neuron pairs with the highest dimorphism in synaptic and functional connectivity

To identify the neuron pairs with the highest dimorphism in synaptic connectivity, we analyzed the combined synaptic weights as described above. Neuron pairs where the combined SEM was greater than or equal to 5 were excluded due to lack of consensus between the available connectomes. The remaining pairs were sorted based on the measured dimorphism defined in the previous section. The top 36 neuron pairs identified through this method are shown in S5. For visualization purposes, we excluded redundant neurons to ensure that each neuron class appeared only once.

To identify the neuron pairs with the highest dimorphism in functional connectivity, we excluded neuron pairs where the *p*-value of the difference between hermaphrodites and males was greater than or equal to 0.01. Thereafter, we sorted functional dimorphisms using the absolute difference between their average values in hermaphrodites and males. The top 36 neuron pairs identified through this method are shown in S5. To match their synaptic counterparts, sensory-sensory neuron pairs and stimulus-neuron pairs were excluded. Pairs not measured in the connectome data (e.g., the pharyngeal neurons and ventral nerve cord motor neurons) were also excluded. For visualization purposes, we again excluded redundant neurons to ensure that each neuron class appeared only once.

## Acknowledgments

We thank Mark Alkema, Javier Apfeld, Grace Comer, Michael Francis, Oliver Hobert, Jinmahn Kim, Albert Lin, Meital Oren-Suissa, Doug Portman, and Paul Sternberg for helpful discussions and suggestions.

## Author Contributions

Conceptualization: M.S., E.Y., and V.V.; Methodology: M.S., M.T., E.Y., and V.V.; Microscope design and build: M.T. and V.V.; Microfluidic setup design and development: M.S. and L.L.; Reagent provision (pheromone): F.C.S; Calcium imaging experiments: M.S.; Neuron identification (NeuroPAL): X.F. and E.Y.; Calcium data analysis and visualization: M.S., M.T., S.R.; Drop test assays: M.S. S.L.; Synaptic data provision: S.C., C.K.; Website and data availability: M.S. and S.R.; Writing - Original draft preparation: M.S.; Writing - review and editing: E.Y. and V.V.; Funding acquisition: E.Y. and V.V.

## Supplementary figures

**Figure S1.**
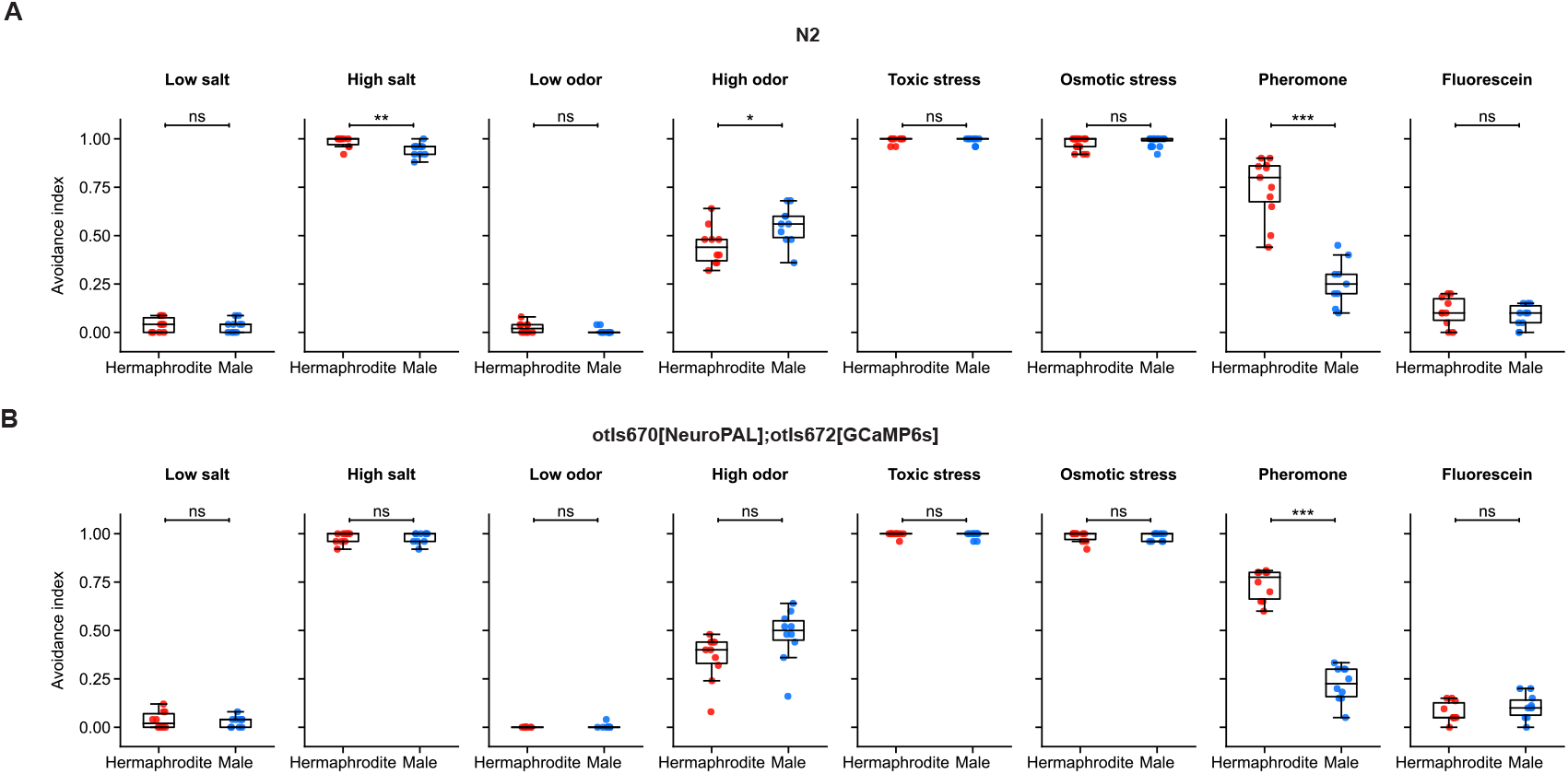
(supplement for Fig. 1C) Behavioral responses to stimuli. Avoidance index in **A**. wild-type (N2) animals and **B**. NeuroPAL GCaMP6s (OH16230 - otIs670[NeuroPAL];otIs672[GCaMP6s]) animals.

**Figure S2.**
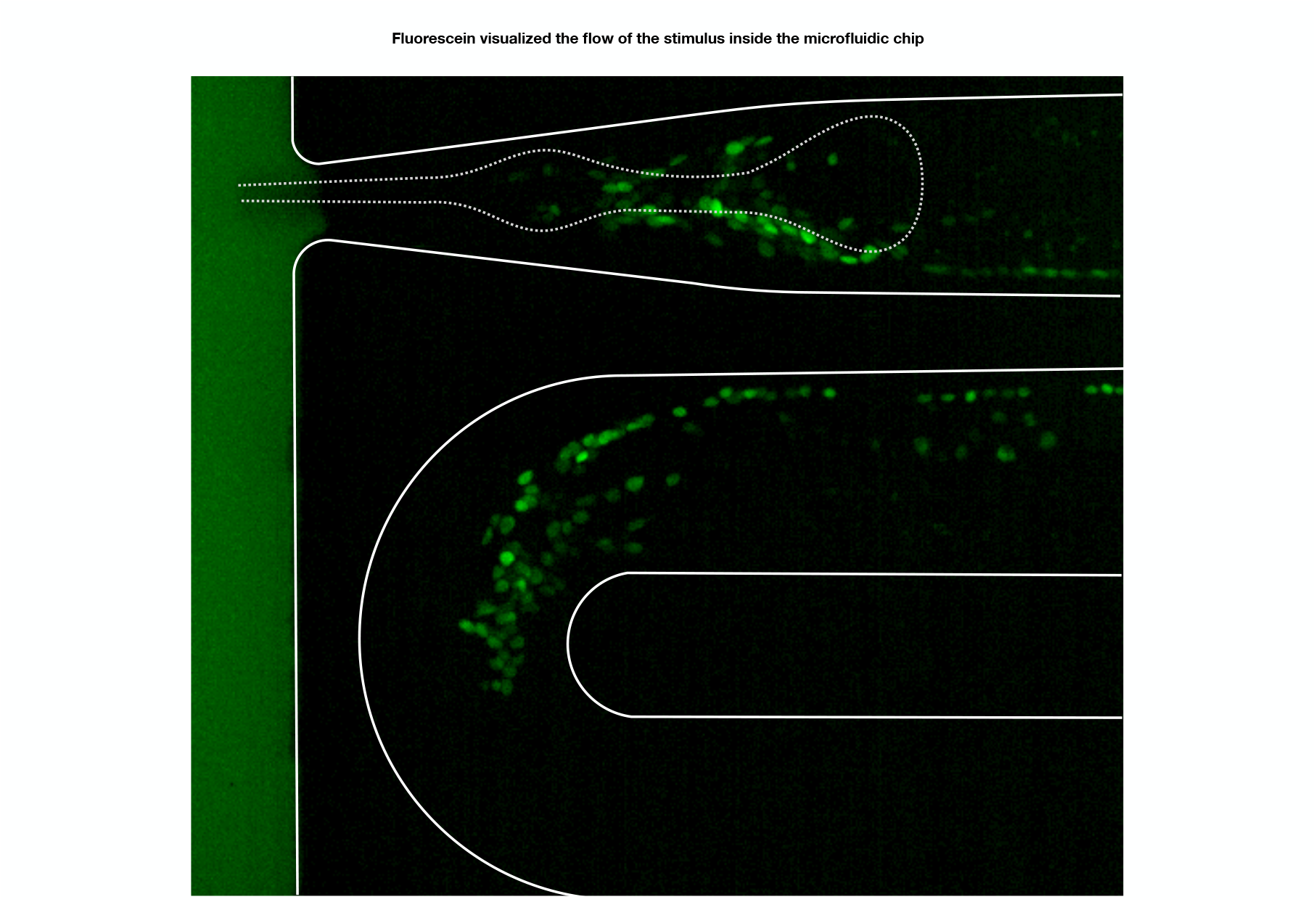
(supplement for Fig. 1E) Using fluorescein to visualize stimulus flow. Fluorescein is used to ensure proper stimulus flow across the animal’s nose, with a tight seal (blocking fluorescein from entering the body chamber) and no air bubbles (these obstruct the flow).

**Figure S3.**
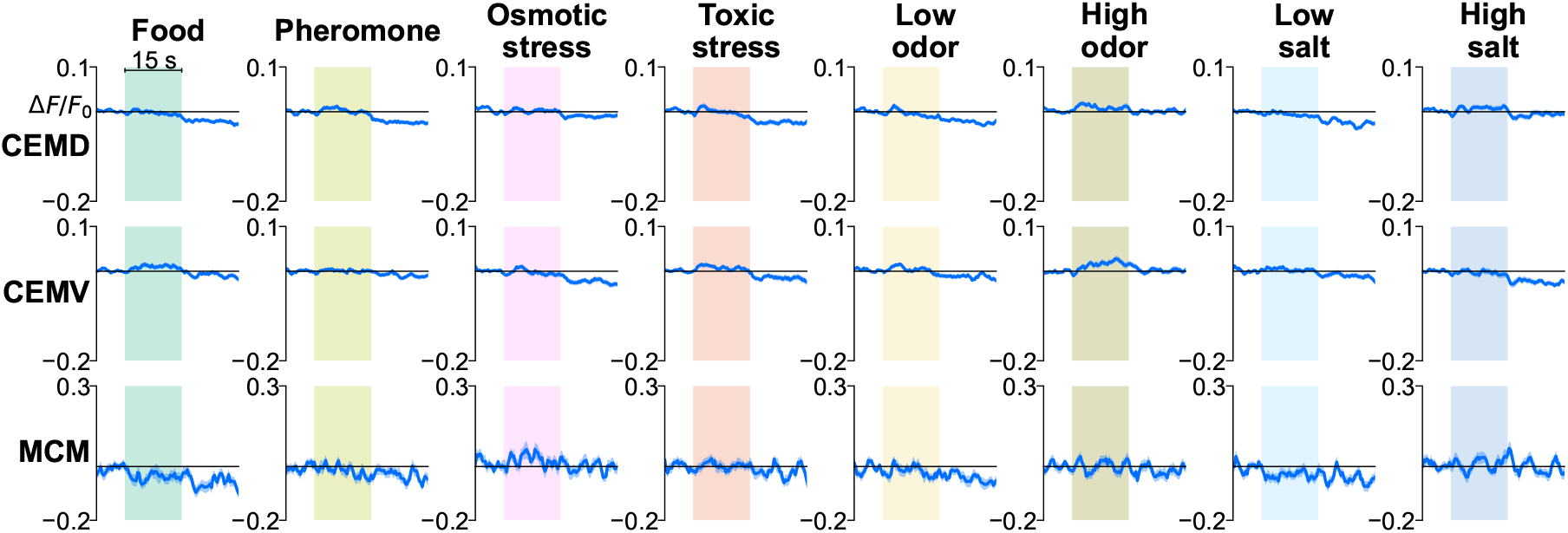
(supplement for Fig. 2B) Male-specific head neurons were unresponsive to stimuli. The CEM and MCM neuron classes were unresponsive to the stimuli.

**Figure S4.**
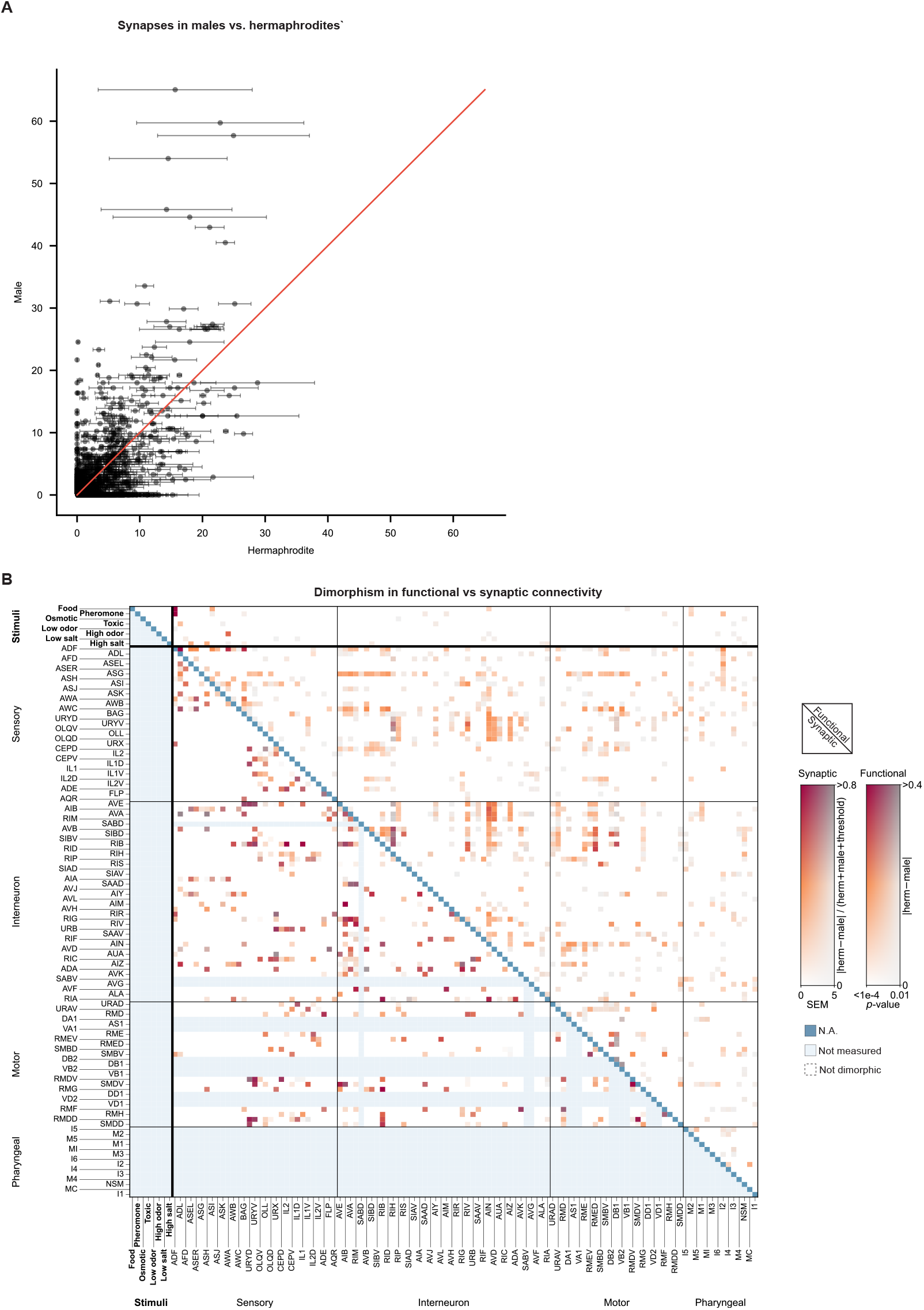
(supplement for Fig. 4) Connectivity comparisons. **A**. Relationship between hermaphrodite and male synapse counts for all pairs of neurons. Error bars represent the variation of synapses seen across multiple hermaphrodite datasets. **B**. Larger version of Fig. 4E with neuron names added.

**Figure S5.**
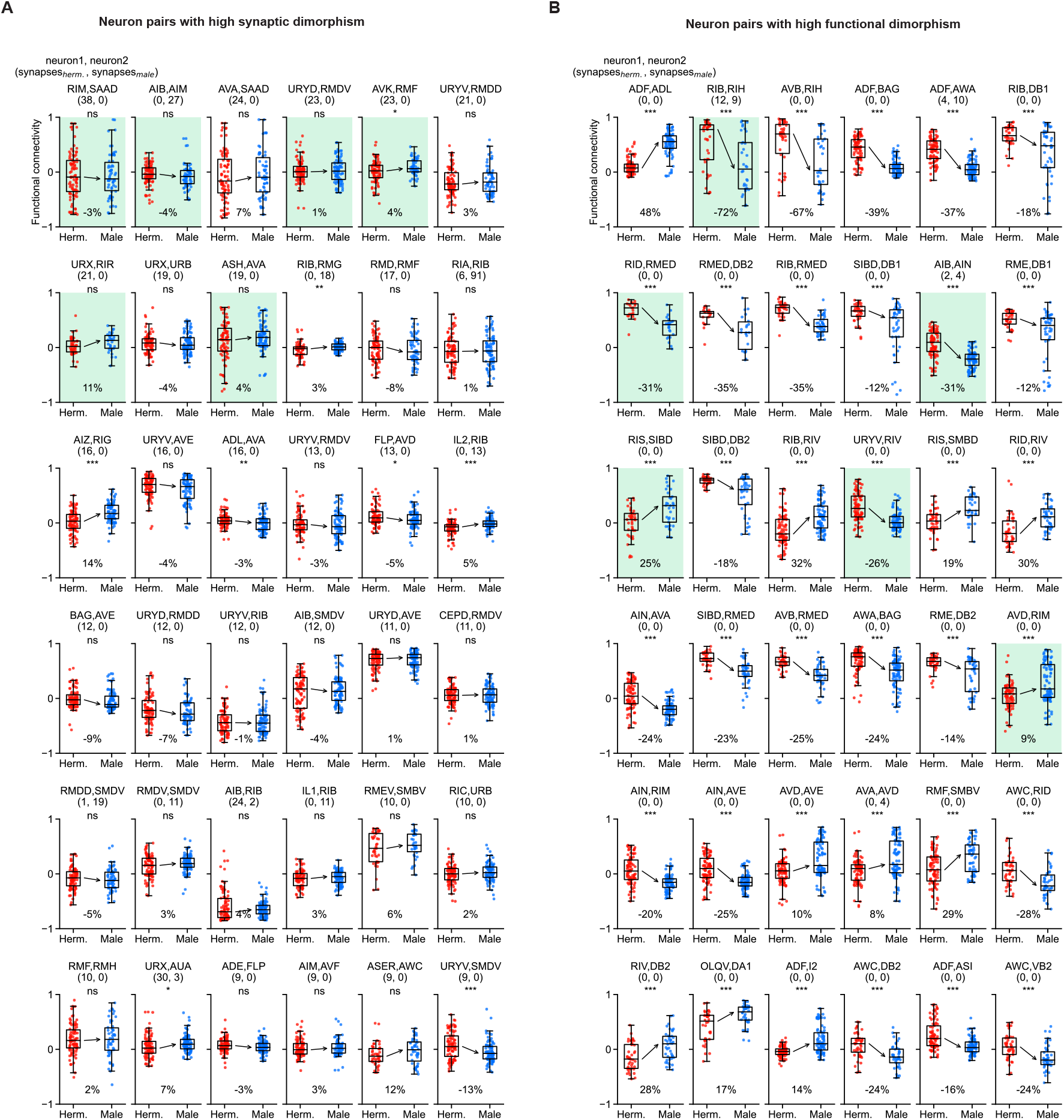
(Supplement for Fig. 4FG) Sexually-dimorphic connectivity. **A**. The 36 neuron pairs with highest synaptic dimorphism. The pairs chosen for Fig. 4F are highlighted (these exclude redundant representation of neurons and sensory-sensory neuron pairs). **B**. The 36 neuron pairs with highest functional dimorphism. The pairs chosen for Fig. 4G are highlighted (these exclude redundant representation of neurons and sensory-sensory neuron pairs).

**Figure S6.**
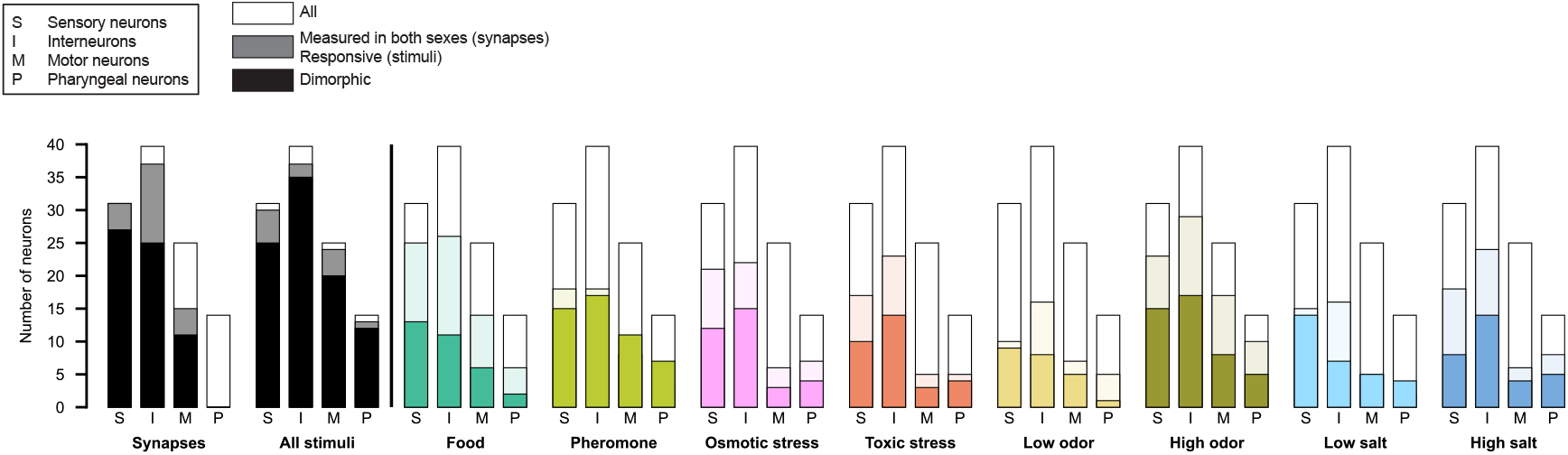
(Supplement for Fig. 5B) Responsive and dimorphic neurons classes. Responsive and dimorphic neurons using the non-canonical subclasses shown in Figs. 3 and 4).

